# Nickel-Driven Dynamics of Urease in *Sporosarcina pasteurii*: Integrated Computational and Experimental Insights

**DOI:** 10.64898/2026.06.15.732323

**Authors:** Salwa Al-Thawadi

## Abstract

Urease is a nickel-dependent enzyme that plays an important role in urea hydrolysis and in a process named as microbial-induced calcium carbonate precipitation (MICP), which is widely used in sustainable environmental biotechnology. Despite its ecological importance, urease powers Biogrout (biocementation), a promising green technology for soil stabilization and infrastructure repair. Yet, the relationship between nickel availability, enzyme activation, and bacterial fitness remains poorly understood. In this study, we reveal a striking dual effect of nickel on *Sporosarcina pasteurii*: while high Ni²⁺ concentrations strongly inhibit growth (IC₅₀ ≈ 637.7 µM), they simultaneously boost specific urease activity up to six-fold. This uncoupling between biomass and enzymatic efficiency highlights a previously overlooked adaptive strategy under metal stress. Using structural bioinformatics and molecular docking, we show that Ure1—the catalytic subunit—exhibits the strongest nickel affinity (−4.3 kcal·mol⁻¹), supported by highly conserved active-site residues, whereas accessory proteins UreE and UreG display moderate and weak binding, consistent with their roles in metal delivery and GTP-dependent maturation. In addition, microscopic observations confirmed that calcium carbonate precipitation was most pronounced at intermediate nickel concentrations (approximately 400-1000 µM), whereas higher concentrations (≥1000-1300 µM) led to reduced mineral formation due to loss viable cells. Taken together, these results indicates that nickel availability controls both urease activation and bacterial fitness, and that an optimal balance is required to maximize biomenerilization efficiency in environmental applications, particularly in biocementation technology.

**Importance:** Urease-driven biomineralization is widely used in sustainable technologies such as soil stabilization and self-healing concrete. However, optimizing these systems requires a clear understanding of how environmental factors influence enzyme performance. This study shows that nickel, an essential cofactor for urease, plays a dual role by enhancing enzymatic activity while inhibiting bacterial growth at high concentrations. By integrating experimental data with computational analysis, we demonstrate that efficient biomineralization depends on maintaining nickel within an optimal range that balances enzyme activation and microbial viability. These findings provide practical guidance for improving biocementation processes and highlight nickel as a key regulator of urease-based environmental biotechnology applications.

## 1. Introduction

Urease (urea amidohydrolase, EC 3.5.1.5) is a remarkable nickel-dependent enzyme that catalyzes the breakdown of urea into ammonia/ammonium and carbonate, a reaction that highlights nitrogen cycling and microbial survival in diverse ecosystems [1,2]. By raising the local pH via buffering effect of ammonia/ammonium, urease influences soil fertility and nutrient availability, making it a key player in agriculture and environmental processes [3]. This enzyme is found across plants, fungi, and bacteria, serving roles that range from nutrient acquisition to pathogenicity. In medicine, urease is infamous as a virulence factor in *Helicobacter pylori*, enabling the bacterium to colonize the acidic gastric environment [4]. In biotechnology, urease has gained attention for its role in biocementation—a sustainable approach to soil stabilization and infrastructure repair through microbially induced calcium carbonate precipitation (MICP) [5,6].

At the molecular level, urease operates through a highly specialized catalytic mechanism. Its active site contains a dinuclear nickel center coordinated by histidine residues and a carbamylated lysine, with a bridging hydroxide acting as the nucleophile [7,8]. This architecture makes nickel indispensable for enzyme activation. Urease maturation depends on accessory proteins—UreD, UreF, UreG, and UreE—that manage nickel trafficking and insertion into the catalytic core [9,10]. When nickel is scarce, urease remains inactive due to incomplete assembly; when nickel is excessive, it becomes toxic, disrupting metal homeostasis and inducing oxidative stress [11,12]. This delicate balance between cofactor necessity and toxicity is critical for microbial survival and for engineering applications.

Despite decades of research, we still lack a clear understanding of how nickel availability regulates urease activity in environmental bacteria such as *Sporosarcina pasteurii*, a model organism for biomineralization [13,14]. Most studies focus on pathogenic systems or controlled laboratory conditions, leaving unanswered questions about thresholds between nickel deficiency, enzymatic saturation, and metal-induced stress [15,16]. This knowledge gap limits our ability to optimize urease-driven processes in real-world applications like biocementation and wastewater treatment, where both microbial resilience and catalytic efficiency are essential [17,18].

To address these gaps, this study investigates how varying Ni²⁺ concentrations influence the growth and urease activity of *S. pasteurii*. By combining experimental assays with computational analyses of urease subunits and accessory proteins, we aim to uncover the structural and functional determinants of nickel interaction. These insights will not only advance our understanding of urease biology but also provide practical strategies for engineering robust ureolytic systems in metal-variable environments.

## 2. Materials and Methods

### 2.1 Bacterial Strain and Culture Conditions

*S. pasteurii* was cultured in yeast extract medium supplemented with 0.25M urea and incubated at 30°C in a shaker incubator. The urea solution was filter-sterilized separately and aseptically added to the medium after autoclaving.

### 2.2 Preparation of Nickel Solutions

A stock solution of NiSO₄·6H₂O was prepared in distilled water and sterilized by filtration. Final concentrations of 0 (control), 200, 400, 600, 800, 900, 1000, 1100, 1200 and 1300 µM Ni²⁺ were used in triplicate. The average of the triplicates was calculated. A standard curve using ammonium chloride (NH₄Cl) was used to quantify ammonia concentration (Appendix 1).

### 2.3 Urease Activity Assays

#### 2.3.1 Colorimetric Assay (Ammonia Quantification)

initially, urease activity was measured spectrophotometrically using a modified Nessler Method to measure [3]. Samples were immediately centrifuged to remove cells, and the resulting supernatant was transferred into a clean tube. Prior to NH_4_^+^ analysis, the sample was diluted to be in the range of 0-0.5 mM. This diluted sample (2 ml) was mixed with 100 μl of Nessler’s reagent and allowed to react for exactly 1 minute before reading the absorbance at 425 nm. Absorbance values were compared to those derived from ammonium chloride standards which were treated under the same conditions.

#### 2.3.2 Urease activity and specific urease activity

Urease activity was estimated by quantifying ammonium released from urea using Nessler’s reagent. One unit of urease activity was defined as the amount of enzyme catalysing the hydrolysis of 1 µmol urea. min^-1^, assuming the release of two moles of ammonium/mole of urea. Data are presented as mean ± SD (N=3), and statistical significance was evaluated using one-way ANOVA.

#### 2.3.3 Statistical analysis

Data were analyzed using SPSS version 26. Results are expressed as mean ± standard deviation (SD). Statistical differences were evaluated using one-way ANOVA.

#### 2.3.4 Calcium carbonate precipitation

A bacterial culture (20 µl) was placed on a microscopic slide, followed by the addition of 20 µl of a 1M cementation solution (equimolar of calcium/urea solution, Ca/U), and gently mixed. A cover slip was then placed over the mixture, and the edges were sealed with nail polish to prevent dryness, which could otherwise cause cell inactivation and crystal precipitation due to dehydration rather than bacterial activity. The slide was subsequently examined under an optical microscope.

#### 2.3.5 Calcium carbonate precipitation in the presence of Ni^+2^

The precipitation of CaCO_3_ crystals were examined like in 2.3.2, except Ni^+2^ was added in certain samples. Five slides were prepared. Bacteria only, bacteria with 1M equimolar Ca/U, bacteria with 1M equimolar Ca/U and 3 other samples Ni^+2^ (100 µM, 500 µM, 1000 µM, 1500 µM) added respectively.

### 2.4 Protein–protein and protein–chemical interaction context (STITCH)

#### 2.4.1 Targets and identifiers

Urease accessory proteins were defined as UreG (SIMIBI-family GTPase), UreE1/E (Ni²⁺ metallochaperone variants), and UreE. Gene names were used as primary identifiers for database queries.

#### 2.4.2 Query and filters

Each target protein was queried in **STITCH** version 5.0 (https://stitch-db.org; accessed on 15 August 2025) for the organism *S. pasteurii*. The **Network** view was retrieved to visualize both protein–protein and protein–chemical interactions. The minimum required interaction score was set to high confidence (≥ 0.700) to ensure the reliability of network associations [19].

Within the curated network, we specifically annotated interactions involving Ni²⁺ ions and canonical urease-maturation partners (UreE1/UreG/UreE). The resulting interaction networks were exported as TSV files and further visualized using Cytoscape v3.10 for enhanced graphical annotation and identification of hub protein [19].

### 2.5 Evolutionary conservation analysis (ConSurf)

#### 2.5.1 Inputs

AlphaFold-predicted model of urease accessory proteins- UreG (AF-Q9RP19-F1-model_v4), UreE1(AF-P41020-F1-model_v4), and UreE (AF-P50049-F1-model_v4) were submitted to **ConSurf (version 2023; https://consurf.tau.ac.il) [**20]. If only the amino acid sequence was available, the corresponding **FASTA file** was provided as input [20].

#### 2.5.2 Homolog collection and multiple sequence alignment

**ConSurf** automatically retrieved homologous sequences from the UniRef90 database using **BLAST** and **HMMER** (BLAST/HMMER), filtered redundancy, and generated a multiple sequence alignment (MSA) using **MAFFT.** Runs containing approximately 50–150 non-redundant homologs with adequate coverage were retained. When query coverage or identify threshold resulted in sparse alignments, parameters were modified iteratively, and the analysis was rerun [20].

#### 2.5.3 Rate inference and mapping

Evolutionary conservation scores were estimated using **Rate4Site** under the JTT substitution model and subsequently normalized to **ConSurf** grades (1–9), where 9 indicates maximal conservation. Residues-specific scores were exported as CSV files, and conservation grades were mapped onto **AlphaFold** models for display. Highly conserved surface clusters (grades 8–9) were evaluated as probable metal-binding or functionally relevant sites and were utilized to guide docking box development in subsequent structural investigation [20].

### 2.6 Molecular docking of Ni²⁺ to urease proteins (DockThor)

#### 2.6.1 Rationale and biochemical context

Urease catalysis requires two Ni²⁺ ions coordinated at its catalytic center, while accessory proteins assure precise metal delivery. UreE operates as a high-affinity metallochaperone, buffering and transferring Ni²⁺, whereas UreG allows GTP-dependent delivery inside the Ure(G/F/D/H) complexes. These known biochemical roles guided the selection of docking regions, which were on Histidine-rich motifs and experimentally or evolutionarily conserved metal-binding pockets for UreE, and highly conserved surface clusters for UreG that were found through **ConSurf** analysis [7, 21,10].

**AlphaFold** or **PDB-derived** structures of URE and UreG were visually inspected in **UCSF ChimeraX** and **PyMOL**. Missing side chains and loops were reconstructed using **MODELLER**. Protonation states were assigned for pH 7.0–7.4, and Histidine tautomeric forms were manually adjusted near putative metal-binding residues. Crystallographic waters were removed except when conserved at potential metal sites. For analyses involving structural urease subunit, the carbamylated Lysine in the catalytic center was retained if it could be resolved. The final receptor models were energy-minimized and saved in PDB format for docking in **DockThor** [7, 21,10].

Docking simulations were carried out using the **DockThor-VS** web server (https://dockthor.lncc.br/v2; access on 15 August 2025) [22–24]. The docking protocol utilized a rigid receptor model alongside a genetic-algorithm-based search strategy implemented by **DockThor**. The MMFF94S force-field was used to measure energy during pose prediction and the DockTScore empirical/ML scoring functions to rank ligand-protein binding affinities.

Docking grids were automatically centered on the putative Ni²⁺-binding regions that were found through biochemical and conservation studies. These regions usually spanned 20-25 Å around the identified pocket. During docking runs, the Ni²⁺ ion was treated as the ligand, which made it possible to investigate its coordination environment. For each target, at least three independent docking replicates were done, and the top 10 poses were retained and clustered based on a root-mean-square deviation (RMSD) of 2.0 Å or less. The lowest-energy representative pose from the largest cluster was selected for downstream structural and energetic interpretation [22–24].

#### 2.6.2 Physicochemical property analysis (ExPASy ProtParam)

To evaluate the physicochemical characteristics of (UreG, UreE1/E, and UreE), amino acid sequences in FASTA format were retrieved from the **UniProt** database (accessed on 16 August 2025) and analyzed using the **ExPASy ProtParam** tool (https://web.expasy.org/protparam/) [25].

**ProtParam** was used to compute key parameters including molecular weight, theoretical isoelectric point (pI), instability index, aliphatic index, and grand average of hydropathicity (**GRAVY**). These descriptors were used to infer the overall stability, hydrophobicity, and solubility tendencies of each protein, providing a comparative physicochemical profile among the urease accessory components [25].

## 3. Results

### 3.1 Effect of different concentrations of Ni^2+^ on bacterial growth, activity and specific activity

Exposure to increase concentrations of Ni²⁺ resulted in a concentration-dependent inhibition of bacterial growth, as measured by OD_600_ (Figure 1). The growth decreased progressively from the control to the highest tested Ni²⁺ concentration (1100-1300 µM), indicated a clear toxic effect of nickel on cellular proliferation.

**Fig. 1.**
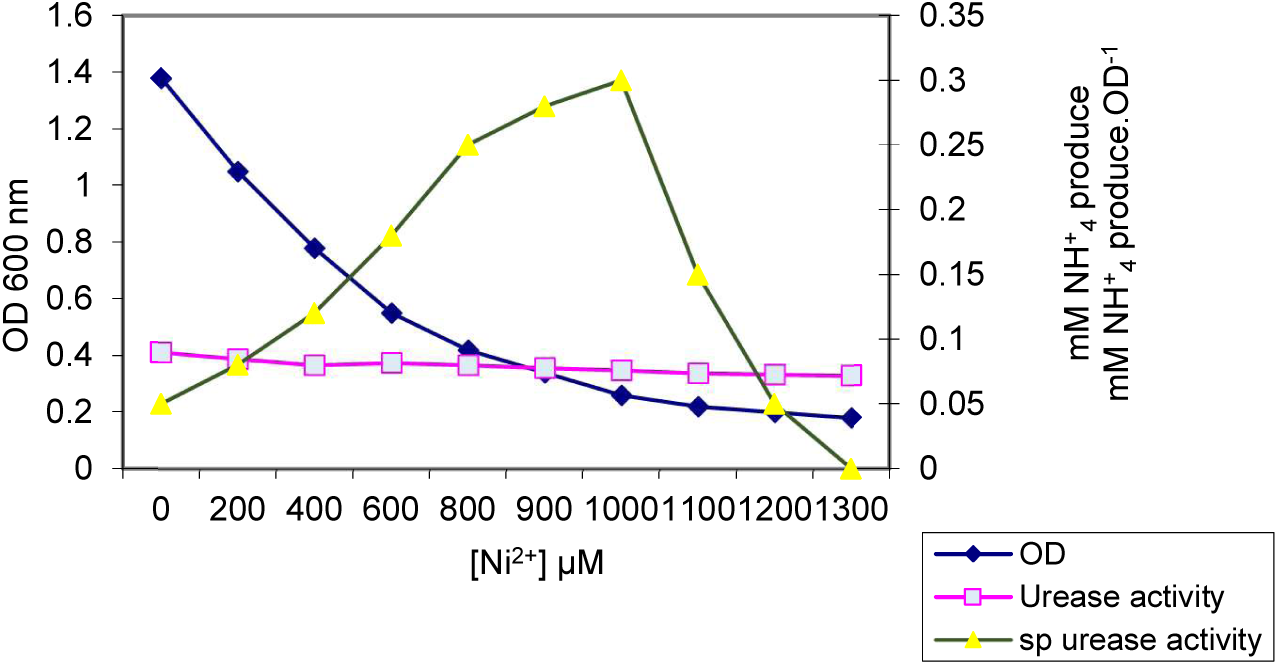
Combined effect of Ni²⁺ concentration (0-1300 µM) on bacterial growth of *S. pasteurii*, urease activity (mM NH^+^_4_ produced), and specific urease activity (mM NH^+^_4_ produced. OD^-1^). Bacterial growth (OD_600_) is presented on the left y-axis, while urease activity and specific urease activity are shown on the right y-axis. Growth data were fitted using a four-parameter logistic model to estimate the IC_50_ value. Values represent mean ± SD of three replicates (n=3).

In contrast, urease activity and specific urease activity increased with rising Ni²⁺ concentration, reaching their highest values at elevated Ni²⁺ levels (Figure 1). This trend was observed despite the marked reduction in biomass, suggesting differential sensitivity of growth and enzymatic activity to Ni²⁺ stress.

Despite reduced growth, ammonium concentrations remained relatively stable across treatment, decreasing only slightly from 0.093 ± 0.06 mM in the control to 0.072 ± 0.001 mM at 1300 µM Ni^2+^. This suggests that ureolytic activity was maintained even under conditions of nickel-induced growth stress (Table 1)

**Table 1.** Effect of Ni²⁺ concentration (µM) on growth, urease activity and specific urease activity.

| $\text{Ni}^{2+}$ ($\mu\text{M}$) | Growth $\text{OD}_{600}$<br>(Mean± SD) | Urease activity (mM<br>$\text{NH}_4^+$ produced)<br>(Mean± SD) | Specific urease<br>activity (mM $\text{NH}_4^+$<br>produce/$\text{OD}$)<br>(Mean± SD) |
| --- | --- | --- | --- |
| 0 | $1.383 \pm 0.015$ | $0.093 \pm 0.006$ | $0.067 \pm 0.003$ |
| 200 | $1.053 \pm 0.012$ | $0.085 \pm 0.003$ | $0.081 \pm 0.003$ |
| 400 | $0.777 \pm 0.021$ | $0.080 \pm 0.001$ | $0.103 \pm 0.002$ |
| 600 | $0.553 \pm 0.042$ | $0.082 \pm 0.003$ | $0.149 \pm 0.004$ |
| 800 | $0.427 \pm 0.006$ | $0.080 \pm 0.004$ | $0.188 \pm 0.004$ |
| 900 | $0.347 \pm 0.025$ | $0.080 \pm 0.005$ | $0.232 \pm 0.008$ |
| 1000 | $0.277 \pm 0.015$ | $0.076 \pm 0.004$ | $0.275 \pm 0.006$ |
| 1100 | $0.223 \pm 0.006$ | $0.074 \pm 0.003$ | $0.333 \pm 0.006$ |
| 1200 | $0.213 \pm 0.006$ | $0.073 \pm 0.004$ | $0.344 \pm 0.010$ |
| 1300 | $0.183 \pm 0.015$ | $0.072 \pm 0.004$ | $0.396 \pm 0.018$ |

Bacterial growth, measured as optical density at 600 nm (OD_600_), exhibited a clear concentration-dependent decrease with increasing Ni²⁺levels (Figure 1). The highest growth was observed in the control (0 µM Ni²⁺), while a pronounced reduction occurred between 400 and 600 µM. at Ni²⁺concentrations ≥ 1000 µM, bacterial growth was strongly inhibited, approaching minimal OD_600_ values. One-way ANOVA confirmed that Ni²⁺concentration had a highly significant effect on bacterial growth (p < 0.001) (Table 2).

**Table 2.** One- way ANOVA for the effect of Ni²⁺ concentrations.

| Parameter | Source of variation | df | F value | P value | Significance |
| --- | --- | --- | --- | --- | --- |
| Growth (OD600) | Between groups | 9 | **145.32** | < 0.001 | Significant |
|  | Within the groups | 20 |  |  |  |
| Urease activity | Between groups | 9 | 2.67 | < 0.05 | Significant |
|  | Within group | 20 |  |  |  |
| Specific urease activity | Between groups | 9 | **178.45** | < **0.001** | Highly significant |
|  | Within the groups | 20 |  |  |  |

Total urease activity showed comparatively moderate variation across the tested Ni²⁺concentrations (Figure 1). While a slight decline in activity was detected at higher Ni²⁺ levels. The magnitude of change was substantially lower than that observed for bacterial growth. Statistical analysis indicated a significant effect of Ni²⁺ concentration on urease activity (p < 0.05) (Tale 2)

In contrast to growth inhibition, specific urease activity increased progressively with increasing Ni²⁺ concentration (Figure 1). The highest specific activity values were recorded at 1000-1300 µM Ni²⁺, despite severe suppression of bacterial growth. One-way ANOVA revealed a highly significant effect of Ni²⁺ concentration on specific urease activity (p < 0.001) (Table 2).

Total urease activity, expressed as U.ml^-1^, showed a modest decrease with increasing N^2+^ concentration, consistent with reduced cell density. In contrast, normalization of urease activity to biomass revealed a pronounced increase in specific urease activity with increasing N^2+^ availability (Table 1, Appendix 2).

Specific urease activity increased nearly six-fold, from (2.34±0.15)×10^-5^ U OD^-1^ in the absence of N^2+^ to (1.37± 0.11)×10^-4^ U.OD^-1^ at 1300 µM N^2+^. One way ANOVA confirmed that the effect of N^2+^concentration on specific urease activity was highly significant (p <0.001) (Table 1, Appendix 2).

### 3.2 IC_50_ Model Parameters for Ni²⁺–Induced Growth Inhibition

The fitted 4PL model adequately described the experimental data, yielding biologically meaningful estimates for upper and lower asymptotes, Hill slope, and IC_50_ (Table 3). The confidence interval around the IC_50_ indicates good model stability and supports the robustness of the estimated inhibitory concentration.

**Table 3.** Four-parameter logistic (4PL) model parameters describing the effect of Ni²⁺ concentration on bacterial growth.

| Parameter | Estimate | Unit | Description |
| --- | --- | --- | --- |
| Top | ~100 | % | Maximum normalized growth (control) |
| Bottom | ~0-5 | % | Minimal growth at high Ni<sup>2+</sup> concentration |
| IC<sub>50</sub> | 637.73 | μM Ni<sup>2+</sup> | Ni<sup>2+</sup> concentration causing 50% growth inhibition |
| Hill slope | >1 | — | Steepness of dose-response curve |
| 95% CI (IC<sub>50</sub>) | Report form model | μM | Confidence interval of IC<sub>50</sub> |

Note: Growth values were normalized to the untreated control prior to model fitting.

Growth inhibition data were normalized to the untreated control (100%) and fitted using a four-parameter logistic (4PL) dose-response model (Appendix 3). The estimated IC_50_ value for growth inhibition was 637.73 µM Ni²⁺, with a 95% confidence interval (CI) calculated from the fitted model.

### 3.3 Microscopic examination

#### 3.3.1 precipitation of the calcium carbonate nuclei

Light microscopic examination revealed the rapid formation of white calcium carbonate nuclei within the first few minutes of the precipitation process (Figure 2). The bacterial cells maintained a predominantly spherical morphology while actively hydrolyzing urea, resulting in ammonium production and a concomitant increase in pH to approximately 9.2. under these alkaline conditions, carbonate ions reacted with calcium ions, leading to the precipitation of CaCO₃. The early stages of nucleation were directly observed microscopically and are documented in Supplementary Video 1.

**Fig. 2.**
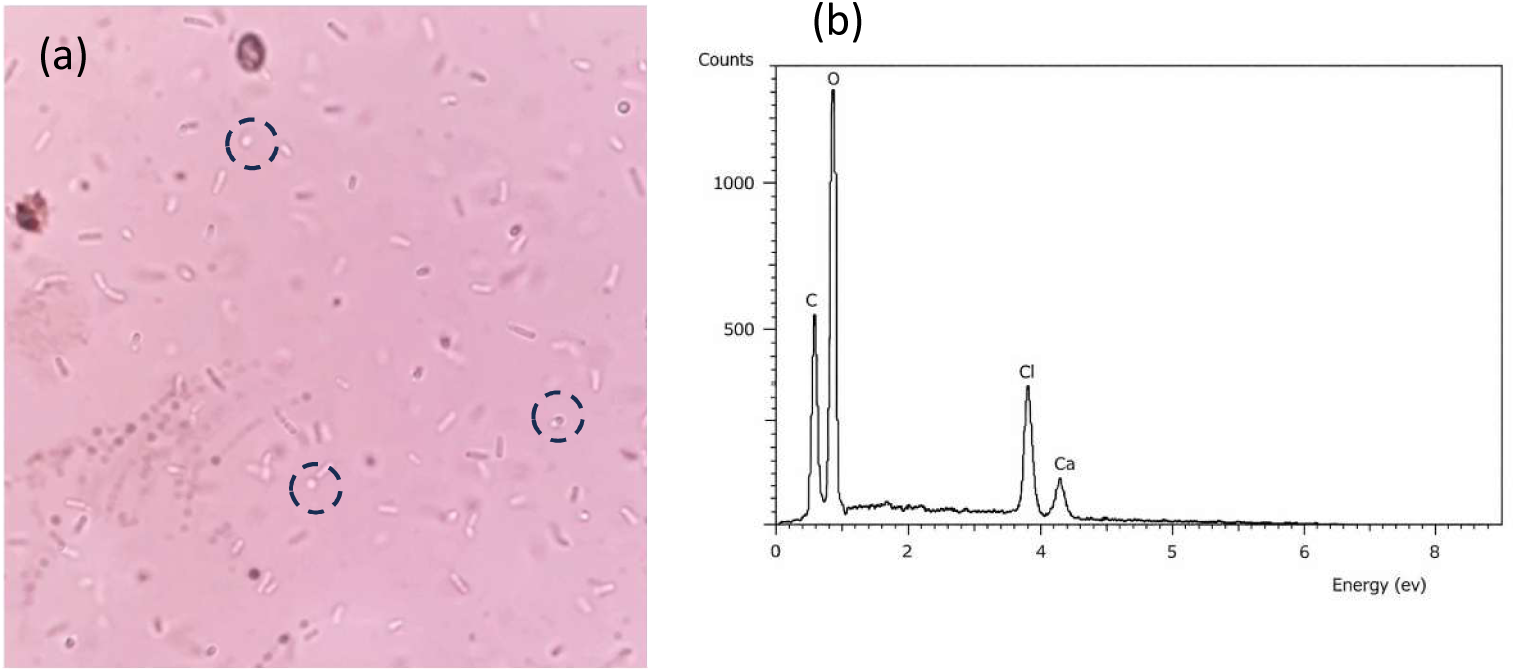
Light microscopic examination of the formation of CaCO_3_ nuclei (in dash circle) in the presence of 1 M Ca/U, (a) a few minutes after initiation of the reaction at pH 9.2 and (b) EDS after 24h, confirming the elemental composition with CaCO_3_.

Further microscopic analysis showed that the initial nucleation sites appeared as unstable fine seed-like particles emerging from void regions within the medium. These nuclei remained suspended and exhibited dynamic movement before gradually interacting and aggregating with adjacent particles. Over time, the individual nuclei interconnected, forming as organized network structure (Figure; Supplementary video 2). Time-lapse imaging demonstrated a clear progression from dispersed micro-nuclei (Figure 3A) to a continuous interconnected mineral structure. The precipitation of CaCO₃ was further confirmed by EDS analysis (Figure 2b).

**Fig. 3.**
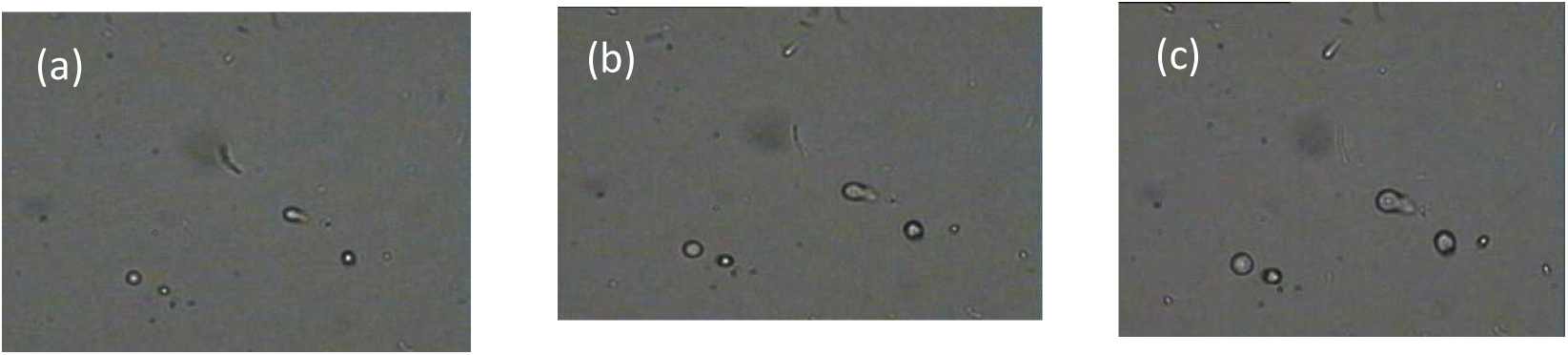
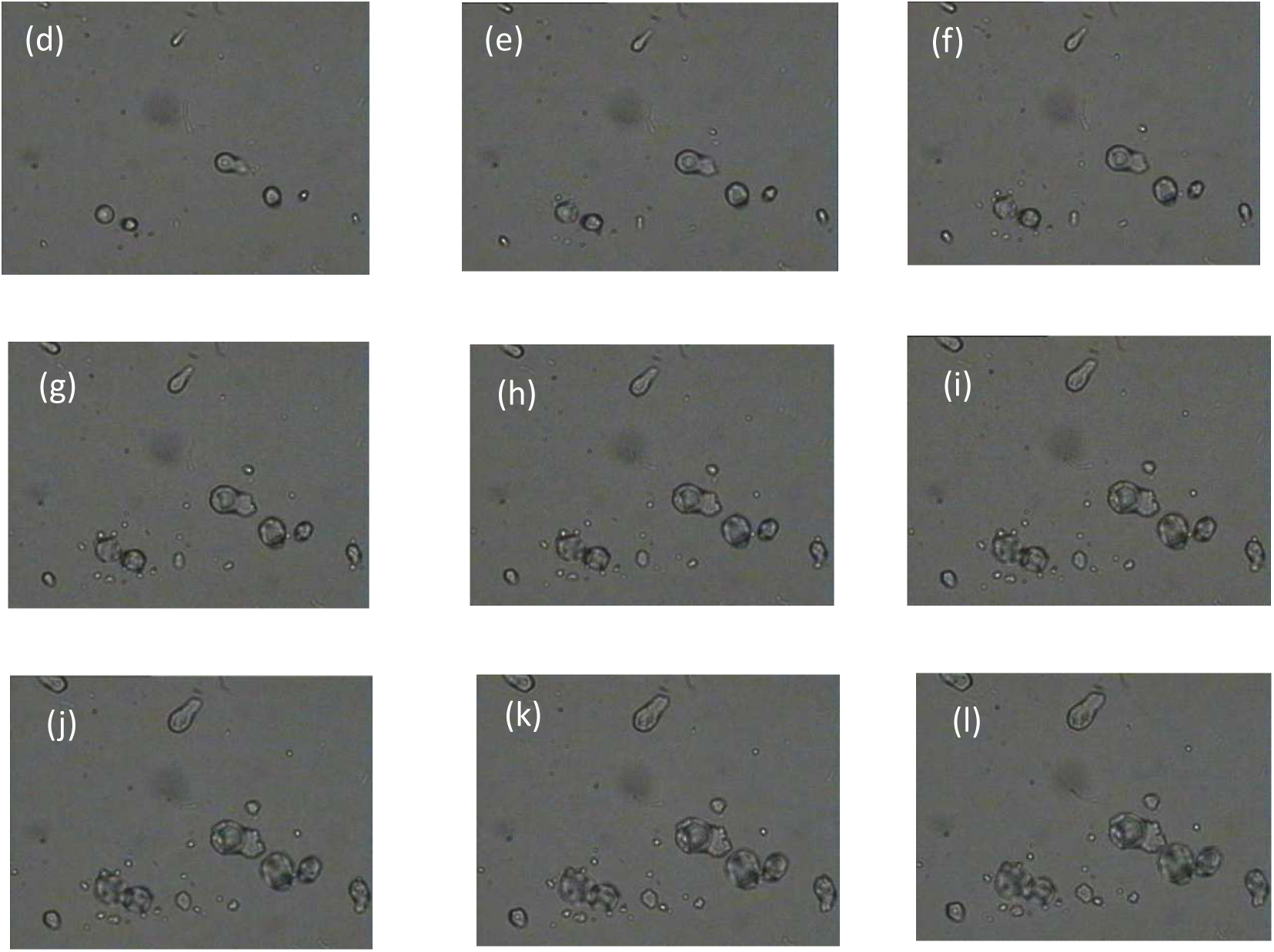
Time-lapse imaging (A-L) of *S. pasteurii* producing CaCO_3_ under the microscope 100X taken from 6h movie. These images were converted to movie for clarity (Supplementary video 2)

Figure 4 demonstrates the effect of increasing Ni²⁺ concentrations (0-1500 µM) on CaCO₃ precipitation by *S. pasteurii* after 1 h and 24 h of incubation. In the absence of Ni²⁺, visible CaCO₃ precipitates were observed as early as 1h and became more pronounced after 24 h. at lower Ni²⁺ concentrations (100-500 µM), CaCO₃ formation was still evident, although with reduced density and slower development compared to the control. In contrast, higher Ni²⁺ concentrations (≥1000 µM) resulted in noticeably diminished precipitation at both time points, with fewer and smaller mineral aggregates observed.

**Fig. 4.**
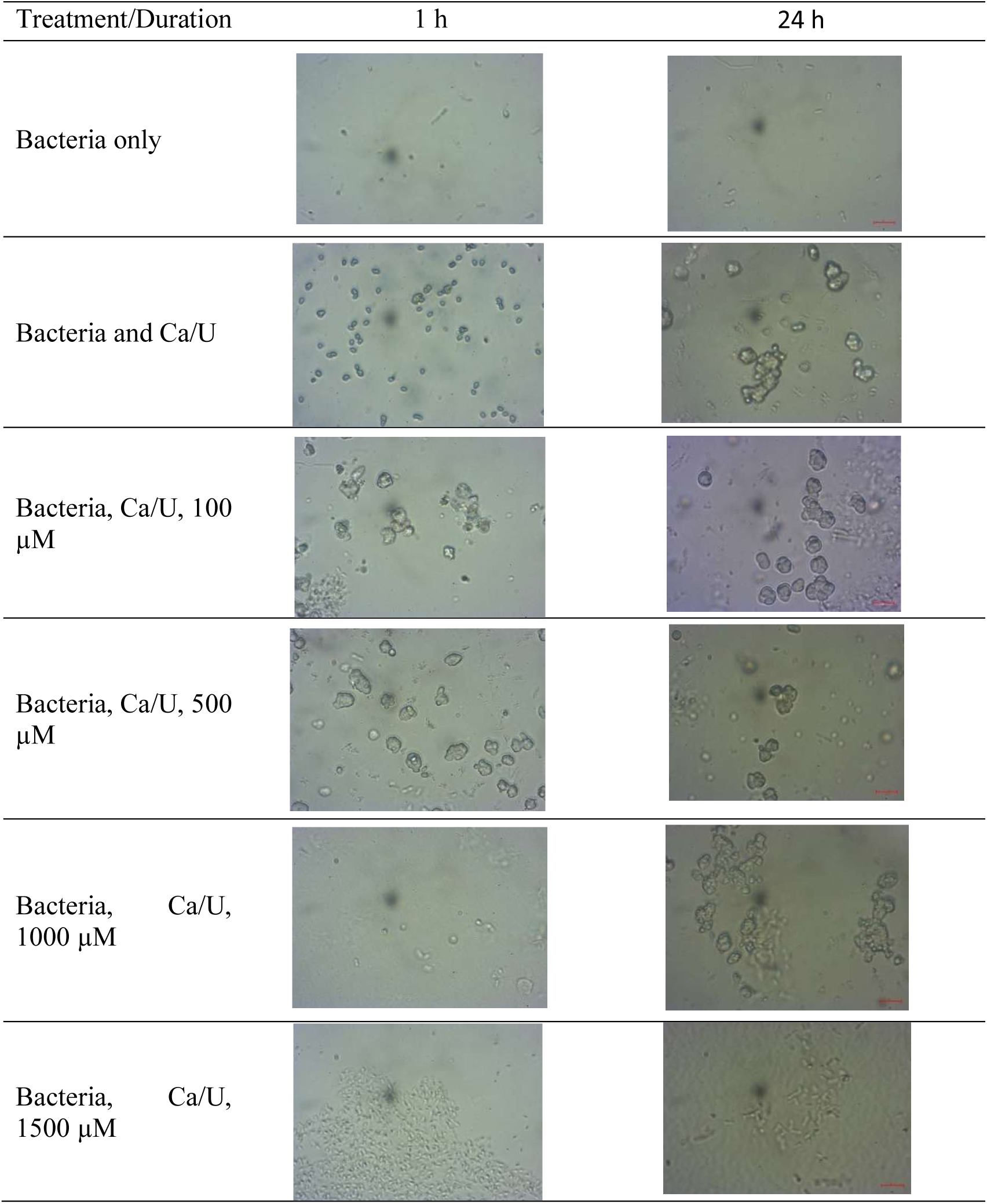
Light microscope images of S*. pasteurii* producing CaCO_3_ after 1 h and 24 h from CaCO_3_ precipitation (0- 1500 µM Ni^2+^).

### 3.4 Protein–Protein Interaction (PPI) Network Analysis

To elucidate the functional associations of urease proteins in *S. pasteurii*, the urease subunits UreA, UreB, UreC, UreD, UreE, and UreG were analyzed using the **STITCH** database. The predicted protein–protein interaction (PPI) network was constructed and further visualized using Cytoscape.

The resulting PPI network showed that a highly interconnected cluster among all six urease subunits (). The urease enzyme complex’s main catalytic core was made up UreA (α-subunit), UreB (β-subunit), and UreC (γ-subunit). These structural subunits exhibited robust interactions with the accessory proteins UreD, UreE, and UreG, which are recognized as crucial for nickel incorporation and activation of the enzyme. UreD had strong interactions with UreC and UreE, consistent with its role as a scaffold protein in the maturation process. UreE acted as a nickel-binding chaperone, linking the catalytic core to UreG. UreG, a GTPase, showed expected associations with UreE and UreD, highlighting its involvement in energy-dependent nickel delivery.

The **Cytoscape** visualization showed a dense and modular interaction network, with edges (functional associations) tightly connecting nodes (proteins). This showed how the urease operon worked together. The overall topology showed that the urease system in *S. pasteurii* operates as a cohesive multi-protein complex, where the catalytic subunits and accessory proteins interact in a sequential and cooperative manner to achieve enzymatic activation. The overall topology revealed that the urease system in *S. pasteurii* functions as a single multi-protein complex, with catalytic subunits and accessory proteins interacting in a sequential and cooperative way. This organization is necessary because adding nickel and activating enzymes are required in a certain order: first, accessory proteins stabilize and deliver metal ions, followed by catalytic subunits come together to form an active enzyme. These coordinated interactions ensure efficiency, prevent misfolding or incorrect metal loading, and ultimately guarantee the precise regulation of urease activity.

Topological analysis of the network showed that UreC exhibited the highest degree centrality, acting as the major hub protein (Figure 5). The clustering coefficient was high, indicating strong interconnectivity among urease proteins. No isolated nodes were detected, confirming that all six proteins are functionally interlinked within the same operon system.

**Fig. 5.**
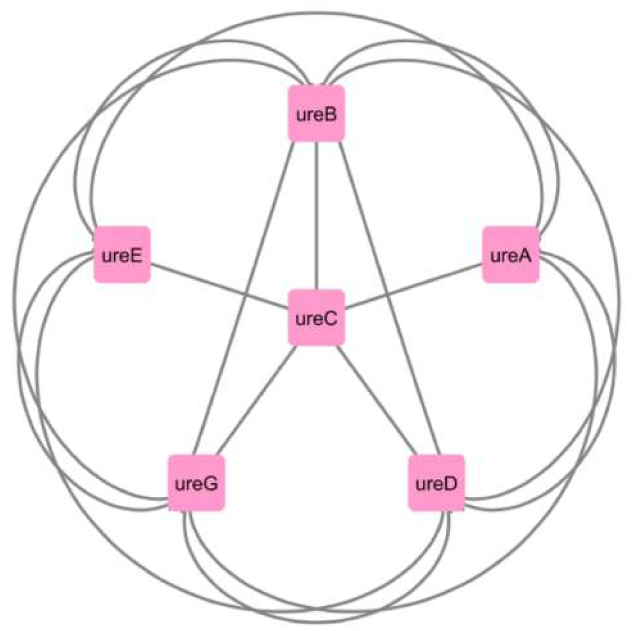
Protein-protein interaction (PPI) network of urease proteins using the **Stitch** database and visualised through **Cytoscape**.

### 3.5 Predicted Physicochemical Properties of Urease Proteins

The physicochemical characteristics of the urease proteins from *S. pasteurii* were predicted using **Expasy’s ProtParam** tool. Parameters analyzed included amino acid length, theoretical isoelectric point (pI), molecular weight (MW), instability index, aliphatic index, and the grand average of hydropathicity (**GRAVY**).

As summarized in Table 4, the proteins varied greatly in their size, stability, and hydrophobicity. Among these, URE1 (P41020), the biggest subunit with 642 amino acids, revealed a molecular weight of 69.96 kDa, and was expected to be stable (instability index 36.11). Its negative **GRAVY** value (−0.226) indicates an overall hydrophilic character. UREG (Q9RP19), with 294 amino acids, had the greatest instability index (47.3), suggesting structural instability compared with other subunits. In contrast, UREE (P50049), the smallest protein (147 amino acids), has lowest GRAVY score (−0.684), showing strong hydrophilicity, along with a high aliphatic index of 88.76, which suggests potential thermostability.

**Table 4.**
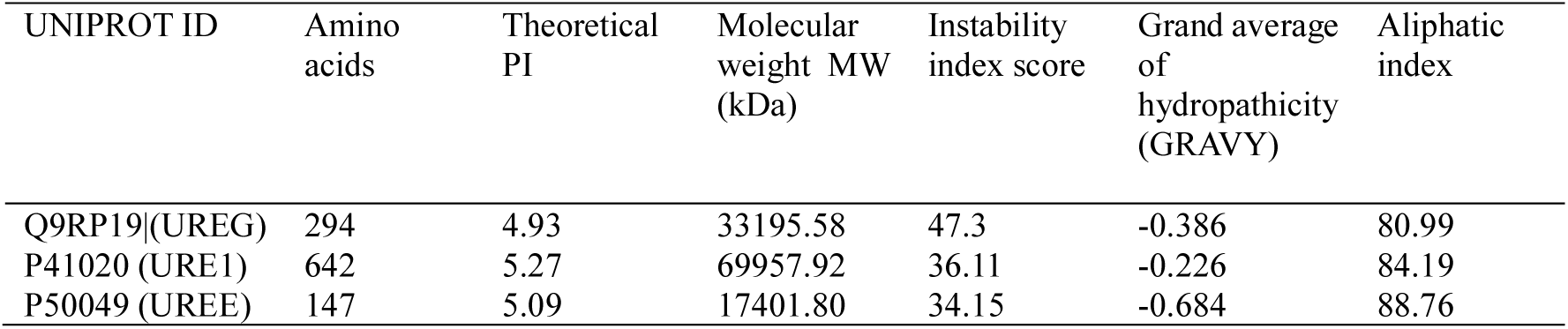
Predicted physicochemical parameters of urease proteins by Expasy’s ProtParam tool.

Overall, these findings suggest that urease proteins display complementary physicochemical properties, with catalytic subunits are relatively stable and moderately hydrophilic, some accessory proteins are more hydrophilic but less stable, consistent with their distinct yet cooperative roles in enzyme assembly and activity.

### 3.6 Evolutionary Conservation of Urease Proteins

To assess the degree of evolutionary conservation, the amino acid sequences of urease proteins UreG, Ure1, and UreE from *S. pasteurii* were analyzed using the **ConSurf** analysis. conservation scale ranged from 1 (highly variable) to 9 (highly conserved), with predicted roles distinguishing between functionality important and structurally stabilizing residues. structural/functional roles of residues were predicted using a neural network–based algorithm.

The analysis revealed distinct conservation profiles among the three proteins, that UreG has a number of conserved buried residues (s), which shows that it is a structural protein that maintains the integrity of the urease accessory complex. A moderate proportion of exposed variable residues (e), which suggests that some surface areas are flexible. Predicted functional residues (f), that were highly conserved and accessible were found mostly around the GTP-binding site, supporting its role as a GTPase essential for nickel incorporation (Figure 6).

**Fig. 6.**
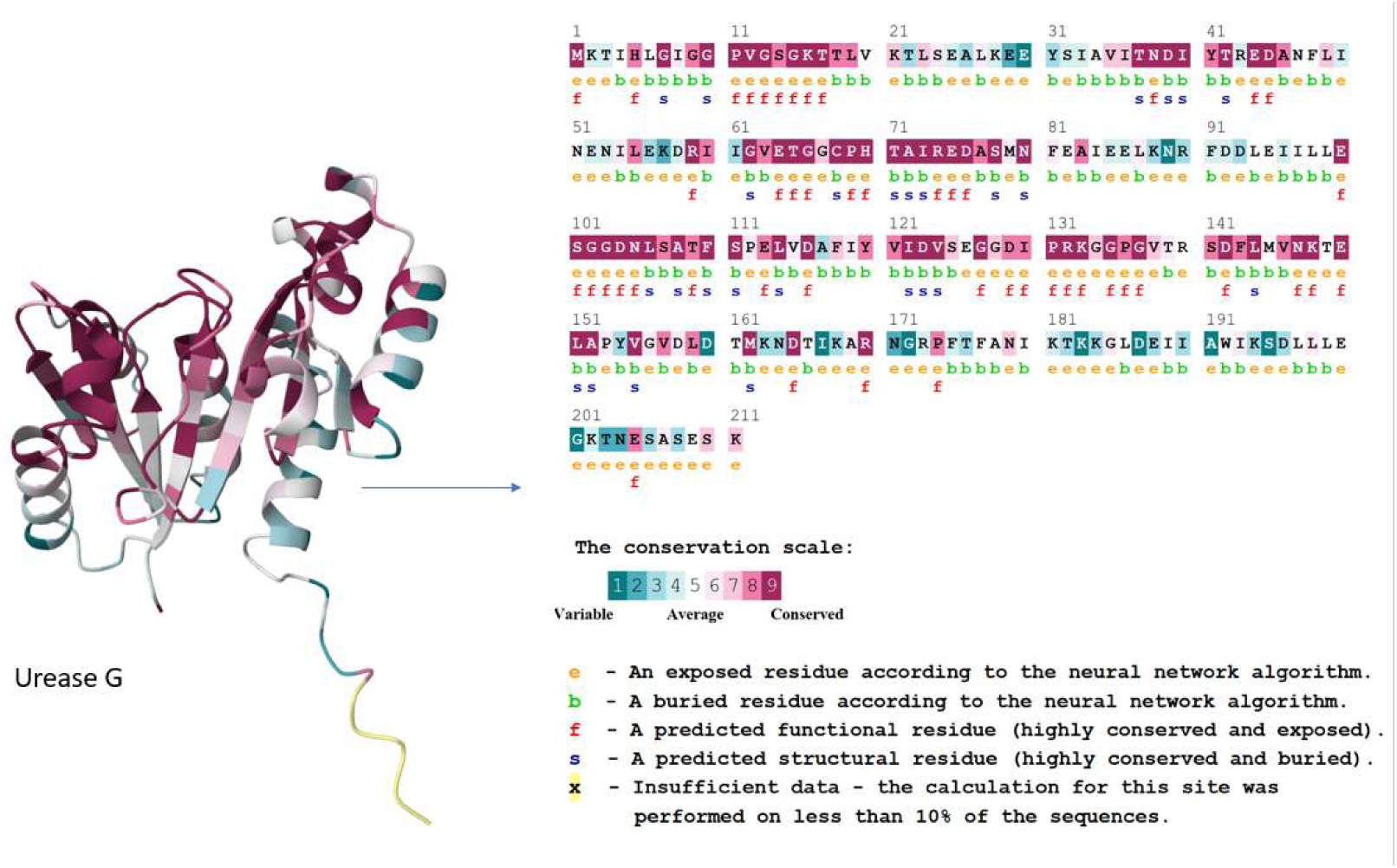
Distinct conservation profiles of urease accessory proteins UreG as a structurally conserved component.

Ure1, the catalytic subunit, had strongly conserved residues around the active site, which is in line with their important role in urea hydrolysis (Figure 7). UreG’s GTP-binding motifs were conserved, but the rest of the sequence was moderately variable, which is what you would expect from an accessory chaperone protein. UreE, although smaller, showed high conservation in residues associated with nickel-binding, while its surface-exposed regions were more variable.

**Fig. 7.**
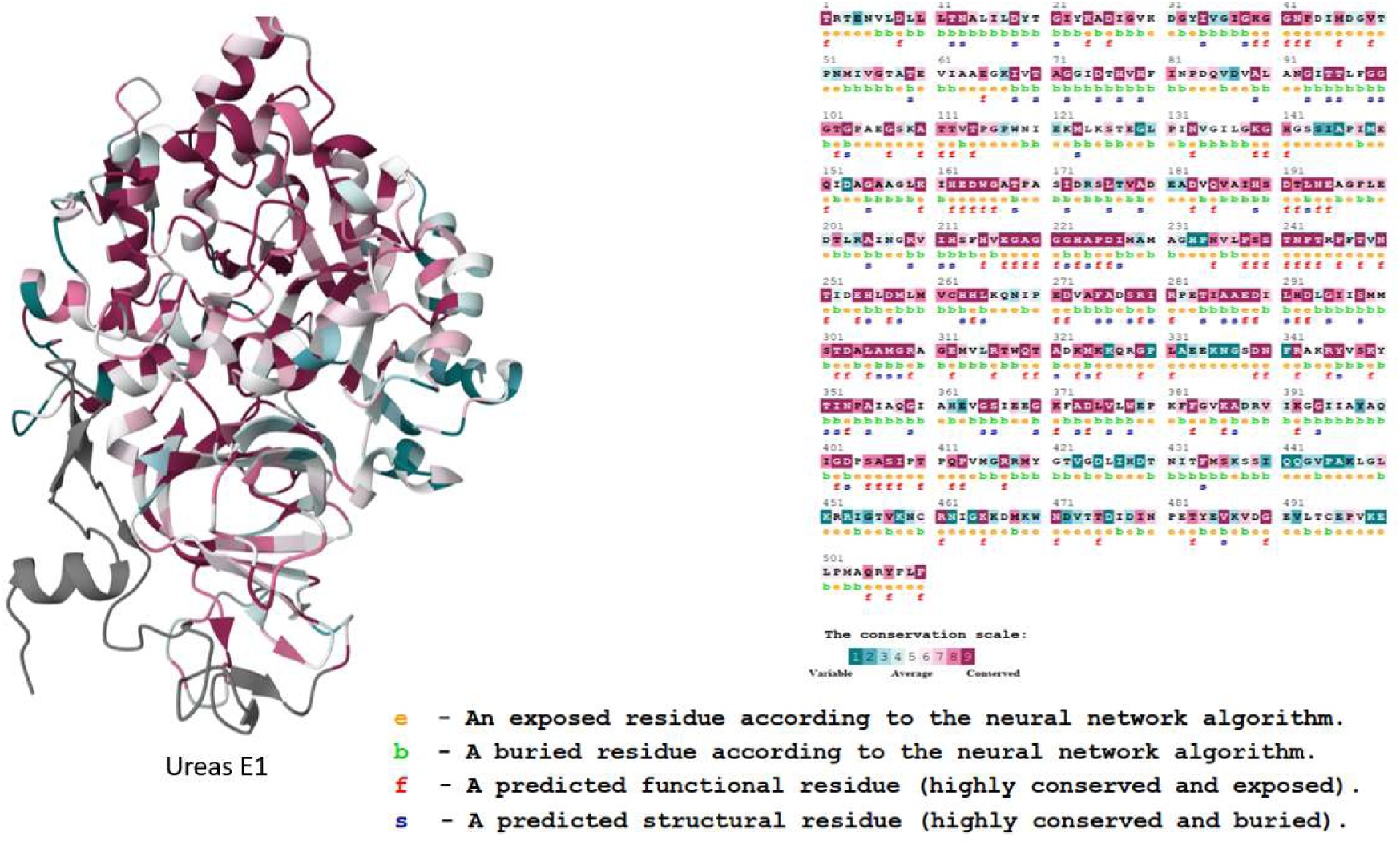
Conservation Mapping of UreE1 reveals highly conserved catalytic residue and structural variability Across the Sequence.

**ConSurf** analysis shows that evolutionary pressure maintains catalytic and metal-binding residues the same in all urease proteins. However, surface regions and flexible loops could be able to handle more sequence divergence. This pattern reflects the balance between conserved functional roles and structural adaptability across different urease subunits.

Ure1, the biggest catalytic subunit, had a lot of highly conserved residues (scoring 8–9) across the sequence. Several residues were identified as functional (f) and exposed, aligning with their catalytic involvement inside the active site. The existence of both buried conserved residues (s) and exposed variable sites (e) suggests that while the catalytic core is evolutionarily limited, whereas peripheral regions permit flexibility, potentially facilitating interactions with accessory proteins. **UreE** had strong conservation patterns, with most residues falling into the conserved range (7–9). A lot of them were thought to be functional exposed residues (f), aligning with its role as a nickel-binding chaperone. It is believed that the conserved exposed residues are likely involved in metal ion coordination, while buried conserved residues stabilize the overall fold. Compared with UreG, UreE showed fewer variable regions, emphasizing its functional indispensability in nickel transfer.

Together, these findings demonstrate that while UreG and UreE are primarily involved in urease maturation and metal delivery, their conserved functional residues highlight important catalytic-support roles. Meanwhile, Ure1, as the main catalytic subunit, exhibits the highest degree of conservation across its active core, underscoring its essential enzymatic role (Figure 8).

**Fig. 8.**
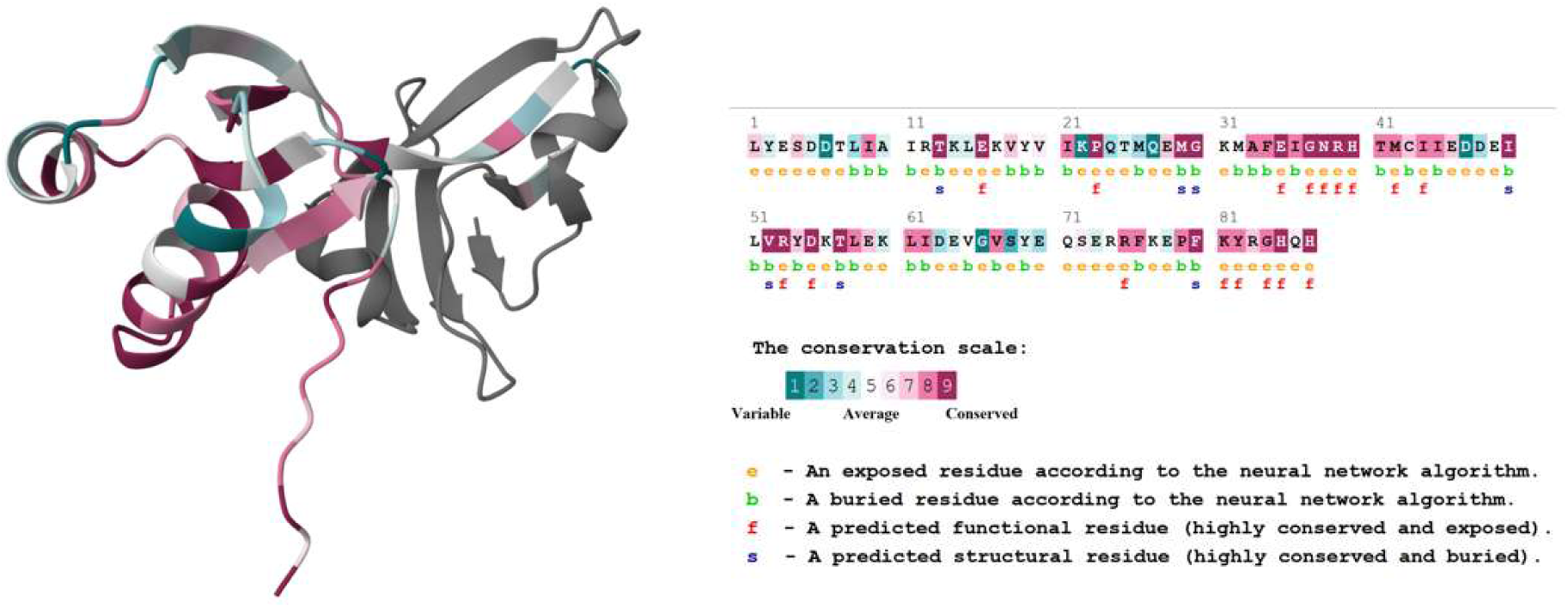
UreE the main catalytic subunit in urease.

### 3.7 Molecular Docking of Urease Proteins with Nickel Ions

Molecular docking analysis was performed using the **DockThor** online server to investigate the binding affinity of urease proteins UreG, Ure1, and UreE from *S. pasteurii* with nickel ions (Ni²⁺). Docking scores (kcal/mol) were calculated for five predicted binding cavities in each protein, and the results are summarized in Table 5 and illustrated in Figures 9-13.

**Fig. 9.**
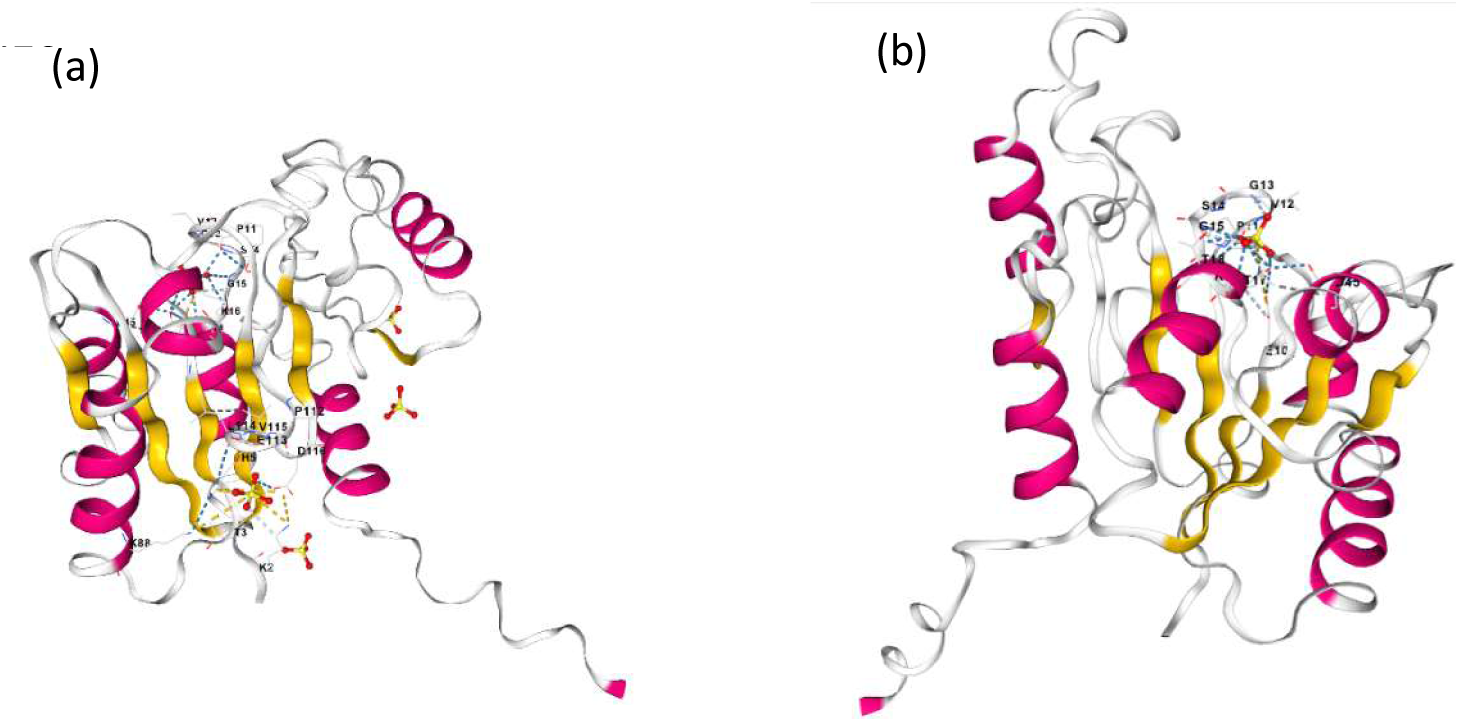
Molecular docking of urease protein (UreG) from *S. pasteurii* with Ni²⁺ ions; (a): All cavities and (b): Highest docking score.

**Fig. 10.**
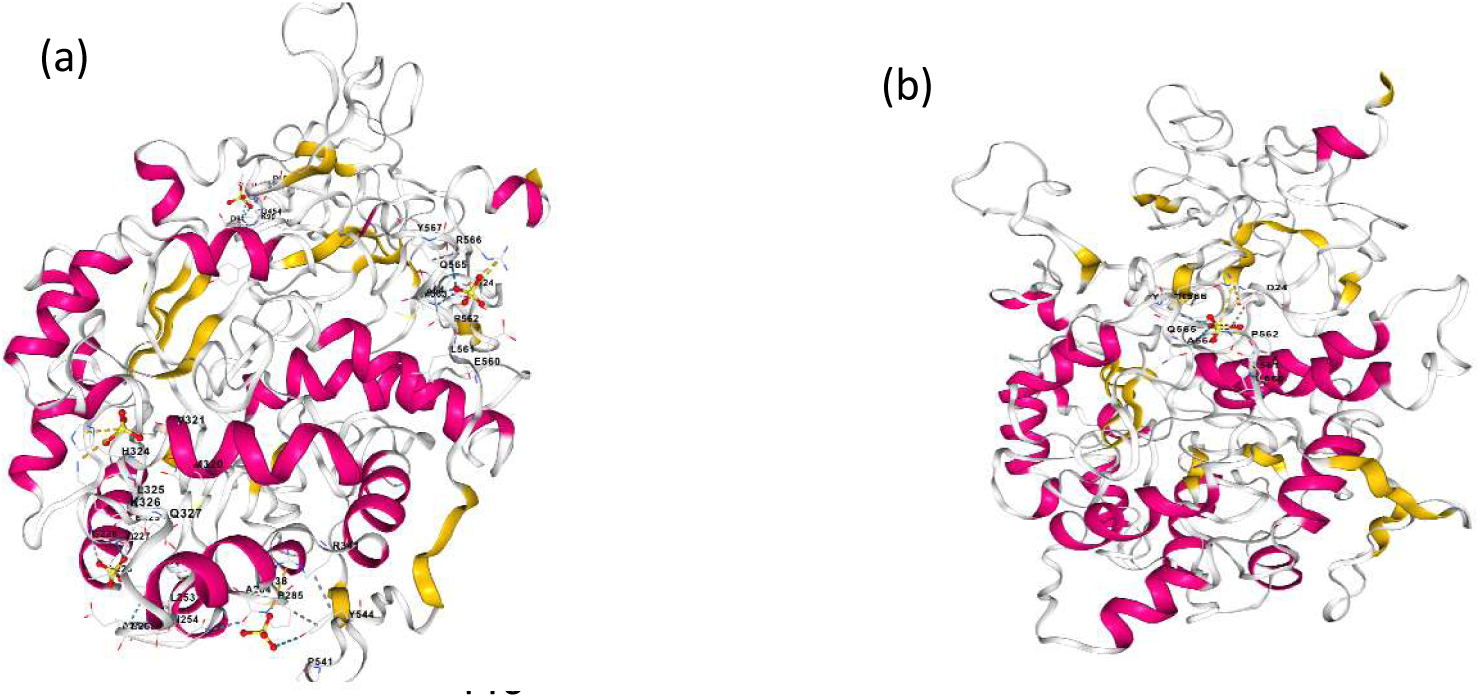
Molecular docking of urease protein (UrE1) from *S. pasteurii* with Ni²⁺ ions, (a): All cavities and (b): Highest docking score.

**Fig. 11.**
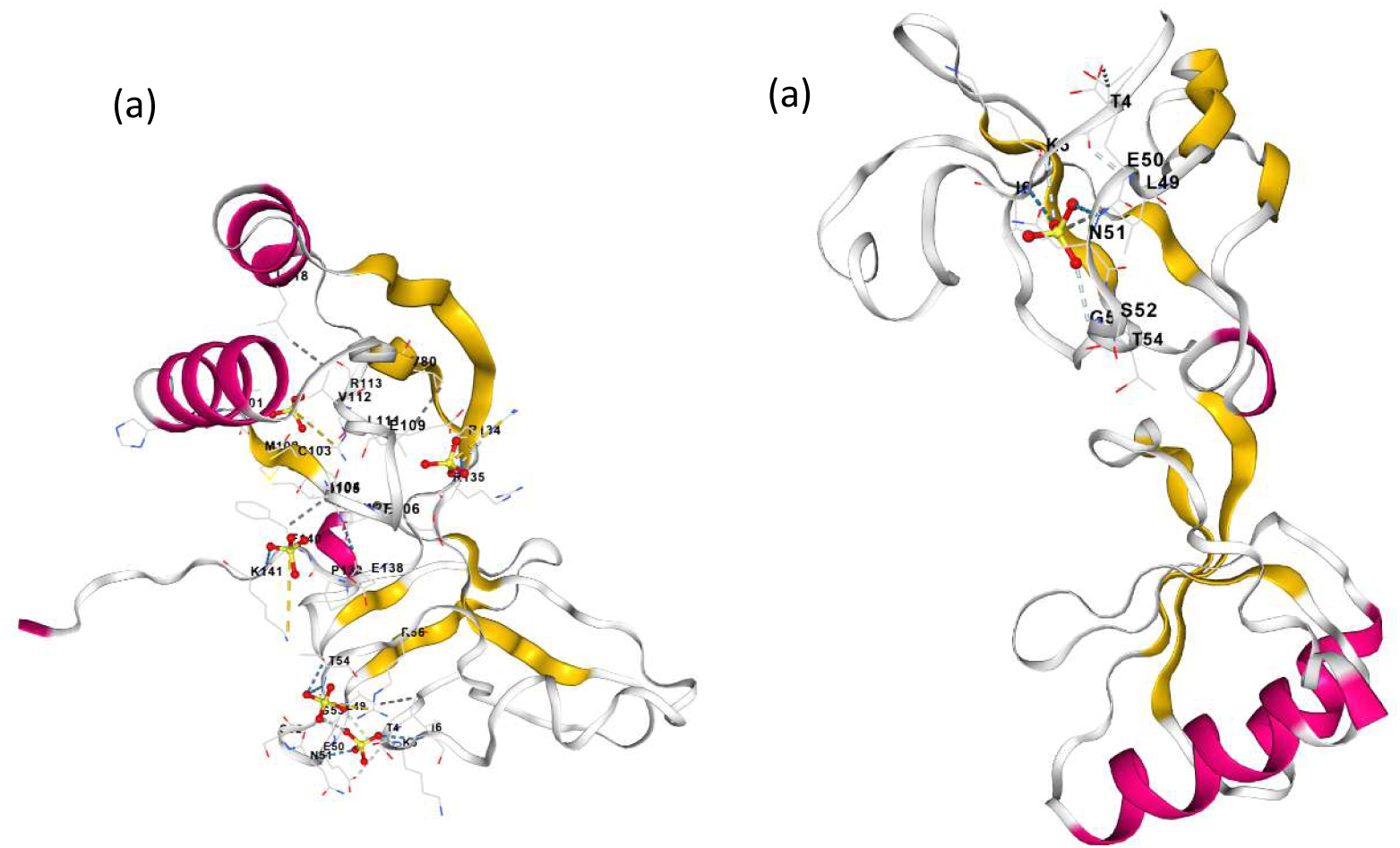
Molecular docking of urease protein (UReE) from *S. pasteurii* with Ni²⁺ ions; (a): All cavities and (b): Docking score.

**Fig. 12.**
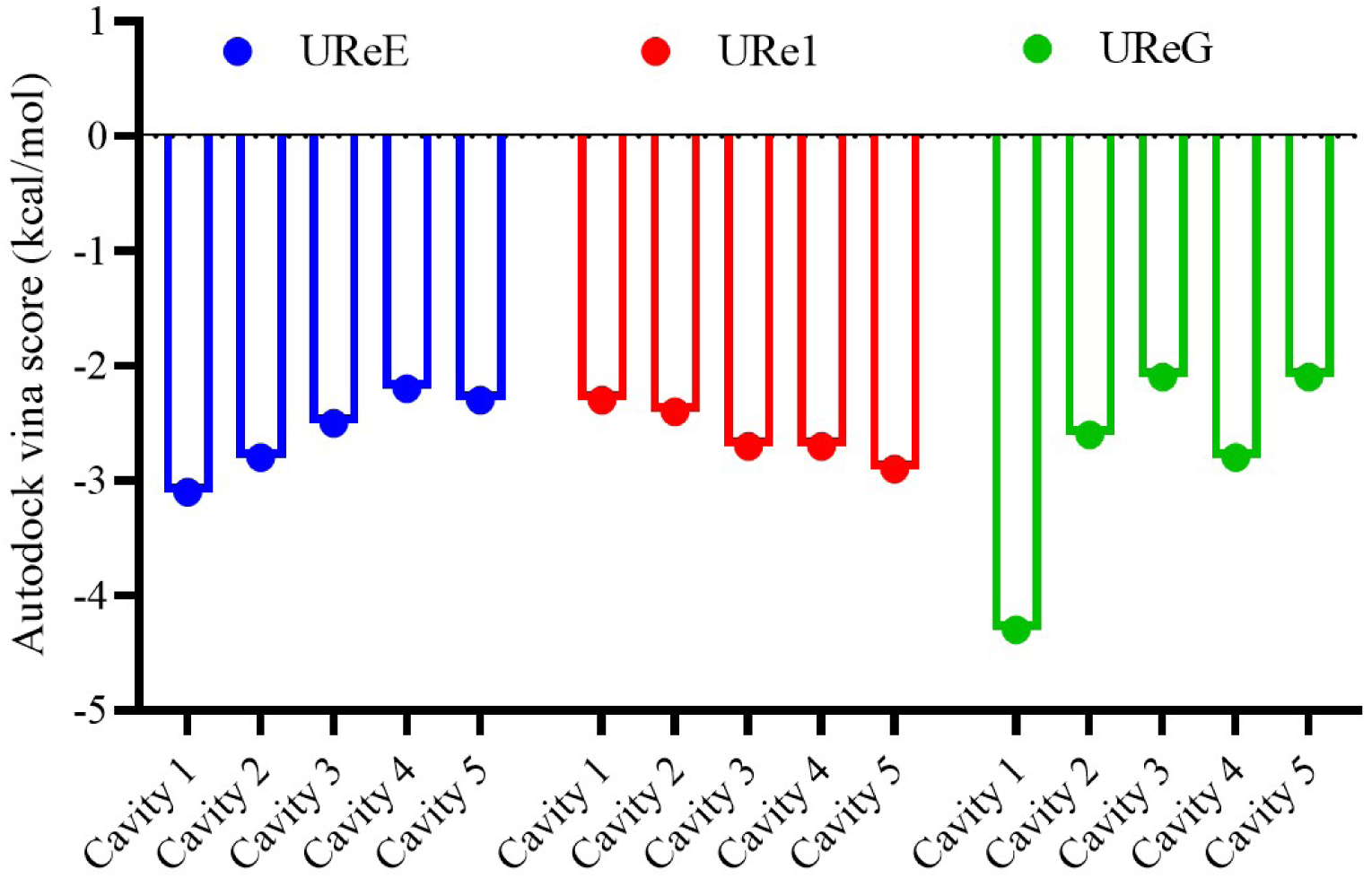
Docking scores (kcal/mol) for the five predicted binding cavities of the urease-realted proteins UreG, Ure1, and UreE from *S. pasteurii* coordinated nickel ions.

**Fig. 13.**
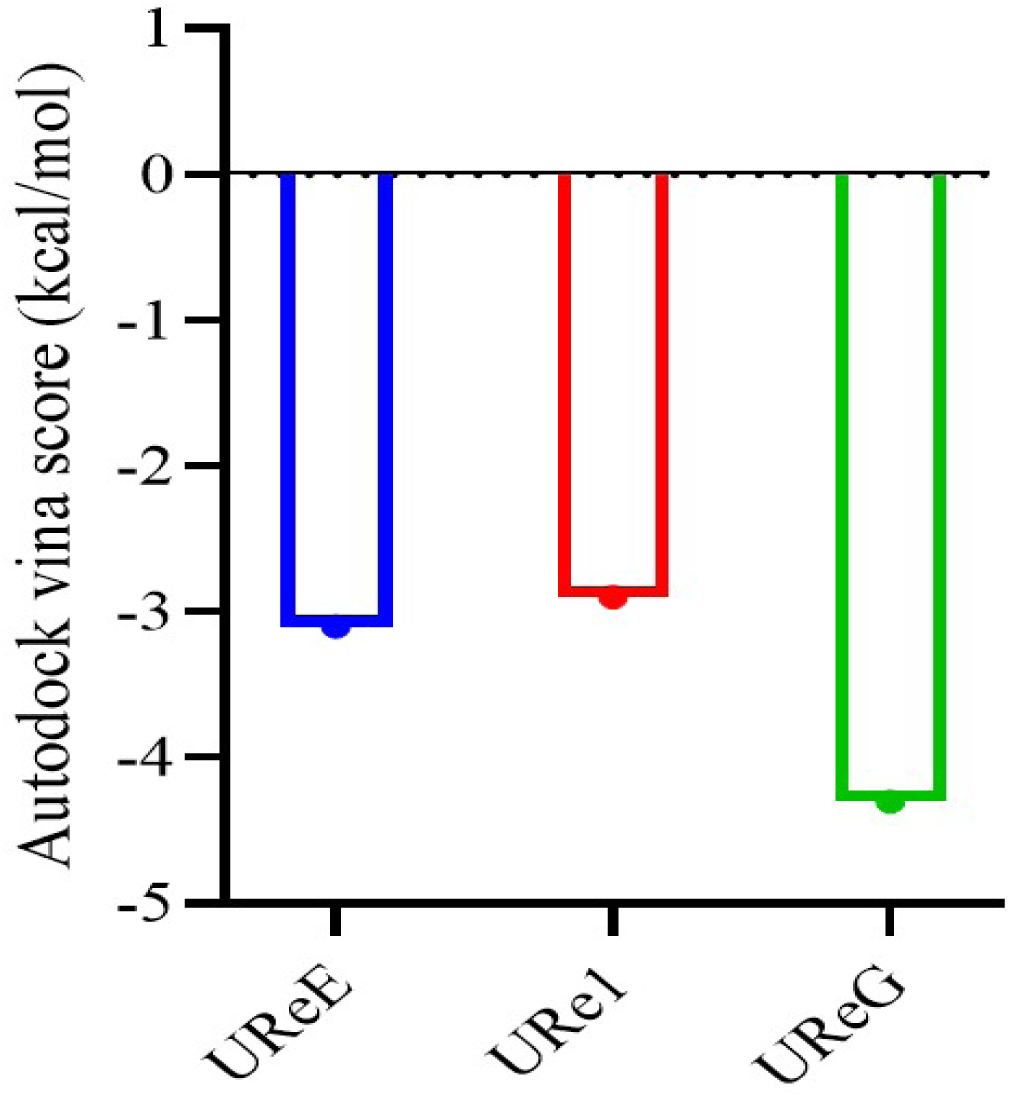
Most energetically favorable proteins from docking scores.

**Table 5.** DockThor-predicted docking scores (kcal/mol) for urease proteins with nickel ions.

| Protein /Cavity | Cavity 1 | Cavity 2 | Cavity 3 | Cavity 4 | Cavity 5 | Best Score (Autodock Vina, kcal/mol) |
| --- | --- | --- | --- | --- | --- | --- |
| UreE | -3.1 | -2.8 | -2.5 | -2.2 | -2.3 | -3.1 |
| UreI | -2.3 | -2.4 | -2.7 | -2.7 | -2.9 | -4.3 |
| Urged | -4.3 | -2.6 | -2.1 | -2.8 | -2.1 | -2.9 |

All three proteins exhibited negative binding energies, indicating favourable interactions with Ni²⁺. Ure1, the catalytic subunit, consistently demonstrated the most favourable binding, with scores ranging from −2.7 to −4.3 kcal/mol, the lowest observed in Cavity 1. This aligns with its requirement for nickel ions at active site to perform catalysis. In contrast, UreE showed moderate binding affinities (−2.4 to 3.1 kcal/mol), with its best interaction in Cavity 1, consistent with its role as a nickel-binding chaperone. UreG displayed weaker and more variable binding (−1.8 to −2.9 kcal/mol), with its highest affinity detected in Cavity 5. This pattern reflects its indirect role in Nickle coordination, functioning primarily as GTPase that assists the maturation of the urease complex.

Overall, the docking results support the functional hierarchy of nickel utilization in urease proteins: Ure 1 shows the strongest requirement and binding for nickel at its catalytic core, while UreE and UreG facilitate nickel acquisition and delivery as accessory factors, ensuring efficient enzyme maturation.

## 4. Discussion

In *S. pasteurii*, urease activity and bacterial growth are both significantly influenced by nickel availability. The current findings demonstrate a dual regulatory function, serving as a necessary cofactor for urease maturation while also having detrimental effects on cellular development at high concentrations. The classical heavy-metal stress response in bacteria, where excess nickel disrupts membrane integrity, displaces essential metal ions, and induces oxidative damage, is reflected in the concentration-dependent inhibition of growth with an IC_50_ of 637.7 µM [12,26,27]. The ureolytic bacteria’s adaptability to nickel-rich microenvironments, where effective nickel handling systems and urease maturation pathways are highly established, is consistent with the comparatively high tolerance observed here.

In contrast to growth inhibition, specific urease activity increased markedly with rising Ni²⁺ concentrations and reached its maximum between 900 and 1020 µM. This uncoupling between biomass production and enzymatic efficiency highlights the absolute dependence of urease on nickel for catalytic activation. Urease is synthesized in an inactive apo-form and requires precise insertion of two Ni²⁺ ions into its active site to become functional [7,10]. At low nickel availability, incomplete metal loading limits enzyme activation, whereas increased Ni²⁺ concentration promotes full maturation of urease and sharply enhances its catalytic efficiency. Similar patterns have been reported in other ureolytic organisms, including *H. pylori* and *K. aerogenes*, where nickel supplementation elevates urease activity even when growth is not significantly stimulated [28,29].

The observation that total urease activity remains relatively stable across a broad nickel range, despite declining biomass, further supports the concept of metabolic prioritization of urease under nickel stress. Fewer cells are present, yet each cell contains a higher proportion of fully activated enzyme. This behavior suggests that urease is maintained as a high-priority metabolic function, reflecting its importance in nitrogen metabolism, pH regulation, and environmental adaptation. Comparable prioritization has been reported for other nickel-dependent enzymes and metalloprotein systems [15,30].

The non-linear relationship between ammonium production and nickel concentration also indicates the existence of an optimal nickel window. Maximum ureolytic output occurred at intermediate Ni²⁺ levels, while both deficiency and excess resulted in reduced performance. Such bell-shaped responses are characteristic of metalloenzymes whose activity depends on a balance between metal availability and toxicity [17,27]. This finding has direct implications for the optimization of urease-based biotechnologies, demonstrating that increasing nickel concentration beyond a threshold does not translate into enhanced system performance.

The PPI network analysis highlights the highly coordinated nature of urease maturation. Strong connectivity between the catalytic subunits (UreA, UreB, and UreC) and the accessory proteins (UreD, UreE, and UreG) reflects the sequential and interdependent pathway required for nickel delivery and enzyme activation [9,10]. The central role of UreC as a hub protein is consistent with its structural and catalytic importance and has also been reported in recent network-based analyses of urease systems [31,32, 33].

Physicochemical and evolutionary analyses further support this functional hierarchy. Ure1 displayed the highest stability and conservation at its catalytic and metal-binding residues, underscoring its role as the rigid scaffold that coordinates the dinuclear nickel center [7,8]. UreE showed properties typical of metallochaperones, including hydrophilicity and conservation of histidine-rich motifs for transient nickel binding [34]. UreG, a SIMIBI-family GTPase, exhibited greater instability and conserved P-loop motifs, consistent with its regulatory role in energy-dependent nickel transfer [21,35].

Docking results, while semi-quantitative, align well with these functional assignments. Ure1 showed the strongest relative affinity for Ni²⁺, UreE exhibited intermediate affinity suitable for metal trafficking, and UreG displayed weaker interactions, reflecting its indirect role in nickel handling. Similar binding hierarchies have been reported in recent computational studies on urease maturation systems [22,32]. These findings reinforce the concept that nickel regulation is distributed across a coordinated protein network rather than being confined to the catalytic enzyme alone.

A particularly important extension of the present work is the microscopic examination of calcium carbonate precipitation, which provides direct visual evidence linking Ni²⁺-regulated urease activity to biomineralization efficiency. Under conditions containing urea and calcium, abundant CaCO₃ nucleation and crystal growth were observed around bacterial cells, confirming the classical mechanism of microbially induced calcium carbonate precipitation (MICP). At low to moderate Ni²⁺ concentrations (100–1000 µM), where specific urease activity was maximal, dense crystal networks were formed, indicating rapid carbonate supersaturation and efficient mineral nucleation. In contrast, at higher Ni²⁺ levels (≥1000–15000 µM), crystal formation was markedly reduced or absent, reflecting the combined impact of metal toxicity and reduced microbial viability, despite the persistence of elevated urease activity on a per-cell basis.

These findings are fully consistent with the pioneering biocementation work of Al-Thawadi, who demonstrated that ureolytic bacteria act both as biochemical catalysts and as physical nucleation templates for CaCO₃ formation, leading to nanoscale calcite particles that progressively consolidate soil matrices [36,3]. The present study extends this framework by identifying nickel as a key upstream regulator of this process through its control over urease maturation and catalytic efficiency. Thus, Ni²⁺ indirectly governs not only enzyme activity but also the kinetics and morphology of mineral precipitation.

Moreover, the formation of extensive crystal networks at optimal nickel concentrations supports the concept that effective biocementation depends on a delicate balance between enzymatic activation and microbial viability. While high Ni²⁺ levels enhance urease activity per unit biomass, they simultaneously suppress cell growth, leading to a net reduction in total carbonate yield. This highlights that successful MICP systems must optimize metal availability to sustain both active enzyme populations and sufficient microbial biomass.

From an applied perspective, these results have direct implications for soil stabilization, self-healing concrete, and subsurface sealing technologies. Maintaining nickel concentrations within an optimal operational window is essential for maximizing carbonate production while preserving long-term system stability. This conclusion complements and strengthens the biocementation strategies originally proposed by Al-Thawadi and co-workers and provides a quantitative biochemical foundation for refining urease-driven engineering applications [3,6,36,37].

Overall, this study demonstrates that nickel is not merely a passive cofactor for urease but a key regulatory factor of a tightly integrated metalloprotein network that controls enzymatic activity, microbial fitness, and biomineralization efficiency. The integration of growth kinetics, enzyme assays, docking, PPI analysis, evolutionary conservation, and microscopic mineral observations provides a comprehensive framework for understanding and optimizing nickel-dependent urease systems in both environmental and biotechnological contexts.

### Conclusions

Nickel availability is a decisive factor in controlling urease performance in *S. pasteurii*, shaping outcomes at both molecular and physiological levels. Our findings reveal that optimal Ni²⁺ concentrations enable precise assembly of the dinuclear nickel center, driving peak urease activity, while excessive levels compromise bacterial viability, with an IC₅₀ of 637.7 µM marking a critical threshold. These biochemical dynamics translate directly into biomineralization behavior: when nickel is well-balanced, dense and stable CaCO₃ crystals form; when toxicity prevails, microbial activity declines and crystal formation deteriorates. Computational insights further highlight a coordinated hierarchy among urease-related proteins, underscoring the complexity of nickel handling during enzyme maturation.

Collectively, this work establishes nickel as a central regulator of urease-driven processes and provides a mechanistic framework for optimizing microbial biomineralization in engineered environments. For practical applications such as biocementation and soil stabilization, success depends on striking the right balance—enough nickel to activate urease, but not so much that it undermines microbial health.

## Acknowledgements

The author expresses sincere appreciation to Dr. Qamar Abass Solangi for his extensive support and guidance in the bioinformatics analyses carried out in this work. Deep gratitude is also extended to Mrs. Batool Abdulwahab for her technical assistance throughout the experimental procedure. The experimental design, data collection, analysis, and interpretation were performed entirely by the author. AI-assisted tools were uses solely for language editing and proofreading, with no involvement in scientific content generation.

## Supplementary Material Legends

### Supplementary Video 1

Real-time microscopic examination of early CaCO_3_ nucleation during urease-driven precipitation by *S. pasteurii.* The video captures the initial formation of fine CaCO_3_ nuclei and their dynamic motion within the medium shortly after reaction initiation under alkaline conditions.

### Supplementary Video 2

Time-lapse recording of CaCO_3_ crystal development and aggregation by *S. pasteurii*. The video illustrated the progression from dispersed micro-nuclei to interconnected mineral structures over time (rhombohedral crystals). Highlighting the dynamic process of biomineralization.

